# Development of a novel protein identification approach to define mitochondrial proteomic signatures in glioblastoma oncogenesis: T98G vs U87MG cell lines model

**DOI:** 10.1101/270942

**Authors:** Leopoldo Gómez-Caudillo, Ángel G. Martínez-Batallar, Ariadna J. Ortega-Lozano, Diana L. Fernández-Coto, Haydee Rosas-Vargas, Fernando Minauro-Sanmiguel, Sergio Encarnación-Guevara

## Abstract

Glioblastoma Multiforme is a cancer type with an important mitochondrial component. Here was used mitochondrial proteome Random Sampling in 2D gels from T98G (oxidative metabolism) and U87MG (glycolytic metabolism) cell lines to obtain and analyze representative spots (regardless of their intensity, size, or difference in abundance between cell lines) by Principal Component Analysis for protein identification. Identified proteins were ordered into specific Protein-Protein Interaction networks, to each cell line, showing mitochondrial processes related to metabolic change, invasion, and metastasis; and other nonmitochondrial processes such as DNA translation, chaperone response, and autophagy in gliomas. T98G and U87MG cell lines were used as glioblastoma transition model; representative proteomic signatures, with the most important biological processes in each cell line, were defined. This pipeline analysis describes the metabolic status of each line and defines clear mitochondria performance differences for distinct glioblastoma stages, introducing a new useful strategy for the understanding of glioblastoma carcinogenesis formation.

**Biological significance:** This study defines the mitochondria as an organelle that follows and senses the carcinogenesis process by an original proteomic approach, a random sampling in 2DE gels to obtain a representative spots sample and analyzing their relative abundance by Principal Components Analysis; to faithfully describe glioblastoma cells biology.

## Introduction

Pediatric solid brain tumors are the most common Central Nervous System neoplasia in childhood and the second most common before 20 years old [1]. In particular, Glioblastoma Multiforme (GbM) or grade IV astrocytoma is the most common and lethal adult malignant brain tumor [2], while in pediatric population GbM occurred only in 8-12% of the population. Nevertheless, in both populations gliomas are characterized by their aggressive medical behavior, a significant amount of morbidity and high mortality rate [3]. GbM is difficult to classify because they diverge considerably in morphology, location, genetic alterations and low consensus among pathologists in their classification [4]. The characterization of gliomas tumors heterogeneity is a priority for the development of better and more precise diagnostic, prognostic and therapy biomarkers.

Mitochondria, the “power house” of the cell, are abundant in brain tissue; its biogenesis, mitophagy, migration, and morphogenesis are crucial in brain development and synaptic pruning. Mitochondria also affect brain susceptibility to injury, play a part in innate immunity, inflammation in response to infection and acute damage, also in antiviral and antibacterial defense [5]. Due to mitochondria play a critical role in numerous bioenergetic, anabolic and cell biochemical pathways [6,7], genetic and metabolic alterations in mitochondria have been suggested to be the cause, or contributing factors, of pathogenesis in a broad range of human diseases, including cancer [8,9]. Several common features of tumor cells can result from mitochondrial deregulation. Furthermore, mitochondria biology support cell transformation during carcinogenesis [10,11], suggesting that its proteome is versatile and that sense the spatial and temporal dynamics of the cell biological processes, from the onset to the end of cancer. Although these advances, the specific role of mitochondria in cancer has not been completely understood, mainly because the huge amount of information about mitochondrial processes in cancer has not been properly integrated.

Despite the utility of proteomics research to get insights into biological processes of cancer disease and knowledge into neuro-oncology, few proteomic studies in gliomas have been performed to date; the few of them are characterized by the elaboration of lists of proteins found to be, either, up or down-regulated in tissue specimens compared to normal brain. This glut of proteomic data generated has been without a unitary approach to establish the feasibility of the existence of key proteins and/or specific signaling pathways regulating cancer development. So far, most of the data generated is lacking coherence, validity, reproducibility and comparability. The problem arises mainly because of the methodological and analytical limitations, and statistical approaches deficiencies. Even more, a lot of the identified proteins in such studies are irrespective of the nature of the background disease [12–14]. Thus, there is the need for proteomic studies in GbM that generate reliable data to be translated into clinical biomarkers, which contribute to improving patient diagnosis and therapies.

To help the understanding of mitochondrial role in the carcinogenesis of GbM, a proteomic signature, related to the biological processes characterizing two stages of cancer disease, was performed by using T98G and U87MG glioblastoma cell lines; which resemble the metabolic transition (Warburg effect) from mitochondrial OXPHOS to glycolysis, as reported during tumorigenesis [15]. Furthermore, a pipeline for functional analysis of differentially expressed proteins in these cell lines was developed. Thus, a Random Sampling (RS) and Principal Component Analysis (PCA), on 2D IEF/SDS-PAGE mitochondrial proteome gels, were performed to evaluate spots abundance and get a representative spots sample for protein identification by MALDI-TOF. Also, PPI networks extension and GOs enrichment analysis were performed to get a metabolism systemic point of view for T98G and U87MG glioblastoma cells. Our results imply that mitochondria are a definitive and unique cancer sensing organelle for cancer development and the elaboration of therapeutic targets.

## Material and Methods

### Cell culture

T98G (ATCC^®^ CRL-1690™) and U87MG (ATCC^®^ HTB-14™) cell lines were maintained in 175 cm^2^ plastic flasks (37°C, 5% CO_2_) in EMEM medium supplemented with 10% fetal bovine serum (FBS). Cells were harvested with trypsin (80-90%) in confluence with trypsin. Washed twice in PBS and used for mitochondria extraction.

### Mitochondria isolation

The mitochondria were isolated by differential centrifugation. Cells were disrupted separately in 250 mM sucrose, 1 mM EGTA, 10 mM HEPES, pH 7.4 at 4°C and centrifuged for 10 min at 1500 × g and 4°C to recover the supernatant. This step was repeated three times. Subsequently, all supernatants were pooled and centrifuged for 10 min at 12000 × g and 4°C to obtain a mitochondrial pellet. The pellets were used immediately or kept at −80°C until use.

### Mitochondrial proteome extraction

T98G and U87MG mitochondrial-associated proteins were obtained according to Hurkman’s protocol modified as follows: Each mitochondrial pellet was resuspended with 500 μl of extraction buffer (0.7 M sucrose, 0.5 M Tris-Base, 0.1 M KCI, 0.03 M HCI, 0.05 M EDTA and 2% β-mercaptoethanol and saturated phenol (500 μl) and incubated for 20 min at −20°C. Then, mitochondrial samples were centrifuged 10 min at 400 × g, 4°C and the phenolic phase was recovered after (12 to 15h at −20°C) 0.1 M ammonium acetate addition. Then, mitochondrial samples were washed twice with ammonium acetate 0.1 M and centrifuged (4000 × g, 10 min, 4°C). Pellets containing mitochondrial proteins were washed with 1 ml of 80% acetone and centrifuged (4000 × g,10 min, 4°C). Supernatants were discarded, and pellets were resuspended in IEF buffer (7 M urea, 2 M thiourea and 0.06 M DTT, 2% ampholytes (3-10 pH) and 4% CHAPS), centrifuged (8000 × g, 30 min 4°C) [16]. Obtained supernatants were recovered and frozen at −80°C until use for 2D electrophoresis.2-DE gels

Each gel (3 T98G and 3 U87MG) was loaded with 500 μg of protein, quantified by Bradford’s method. IEF was performed in acrylamide gel tubes as in [17], briefly gel tubes were prefocused (2500 v, 110uA, 1hr, and 250/hr, per gel), before IEF (125 V, 22 hr). The electrofocused gels were run into a 2D-SDS PAGE (12%) for additional spot separation. 2D gels were fixed and stained with colloidal Coomassie brilliant blue R-250 for image acquisition.

### Image pre-processing

Gels were scanned in a GS-800 densitometer (Bio-Rad, Hercules, CA) and six images were acquired, wrapped and overlapped with PdQuest 8.0.1 software (Bio-Rad). Next, with all six images mixed, a master gel was created by the default PdQuest algorithm from the sum of the intensity of all spots in gel images.

### Random sampling of spots in master gel

To increase the protein capacity to represent and to describe the cellular processes that are carried out in T98G and U87MG cell lines, we randomly selected 400 spots (of 1274 detected by PdQuest) from the master gel, regardless of their size, intensity or abundance difference between cell lines. With the R V3.4 [18] help, a list of 400 random numbers between 1 and 1274 (the number of spots in the master gel) with uniform distribution was generated, which was the number of spots in the master gel. This process ensures that every spot in master gel has an equal chance of being selected and allows to obtain a representative mitochondrial proteome sample [19]. This spot sample was rematched in all gels image to get a more reliable abundance analysis [14].

### Multivariate analysis of spots intensity

To select the spots to be identified, a spreadsheet with the normalized intensity of the 400 spots sampled was exported from PdQuest. The spots abundance was logarithmically transformed and missing values imputed by Random Forest method with the R package RandomForest [20] to perform the multivariate analysis.

The abundance analysis was performed by principal components analysis (PCA) from the correlation matrix of spots intensity with the R package ade4 [21], to get a spot abundance pattern for the cell lines gels [14]. To know if any component could distinguish between the cell lines, the gels score for each component were plotted. Having found the component, with discriminatory capacity, we identified the significant spots in that component with the square cosine of the correlation matrix between the components and the spots. The abundance pattern was obtained by plotting the mean of the logarithm of the intensity of the significant spots between cell lines [22].

### Mass spectrometry

Each selected spot were cut from gel, alkylated, reduced, digested and automatically transferred to MALDI analysis target by a Proteineer SP II and SP robot using the SPcontrol 3.1.48.0 v software (Bruker Daltonics, Bremen, Germany), with the aid of a DP Chemicals 96 gel digestion kit (Bruker Daltonics) and processed in a MALDI-TOF Autoflex (Bruker Daltonics) to obtain the peptide mass fingerprints. One hundred satisfactory shots in 20 short steps were performed, the peak resolution threshold was set at 1,500, the signal/noise ratio of tolerance was 6, and contaminants were not excluded. The spectrum was annotated by the flexAnalysis 1.2 v SD1 Patch 2 (Bruker Daltonics). The search engine MASCOT [23] was used to compare the fingerprints against the SwissProt [24] release 2016 database with the following parameters: Taxon-Human, mass tolerance of up to 200 ppm, one miss-cleavage allowed, and as the fixed modification Carbamidomethyl and oxidation of methionine as the variable modification.

The mass spectrometry proteomics data have been deposited to the ProteomeXchange Consortium via the PRIDE [25] partner repository with the dataset identifier PXD008540.

### Mitochondrial proteins identification

Identified protein gene was tested against MitoMiner database, which stores a collection of genes that encode proteins with strong mitochondrial localization evidence from 56 published large-scale GFP tagging and mass-spectrometry studies [26], to check mitochondrial membership.

### Basic protein-protein interaction (PPI) net construction

Inicial PPI nets were built accords to STRING application [27]. A net was obtained for T98G and another for U87MG with overexpressed and specific proteins in each cell line as bait nodes and adding edges with the following basic settings: Organism > Homo sapiens; meaning of network edges > evidence; active interactions source > Experiments and Databases; minimum required interaction score > medium confidence (0.400); max number of interactions to show, 1st shell > none, 2nd shell none.

### Significative biological process identification

To know the more critical biochemical processes that are taking place in each cell line. First, the initial PPIs were amplified in STRING, with the previous parameters but increasing three times the initial net in the first shell to add proteins and interactions that increase the representativeness of the cellular processes specific to each line. Next, we performed a comparative enrichment analysis based on Gene Ontology [28] of biological processes sets from extended PPI nets. Enrichment was done employing the Cytoscape [29] overrepresentation plugin, Biological Networks Gene Ontology (BiNGO) [30]. As input, we uploaded the UniProt protein identifiers of all the elements in the initial PPI net first and extended net later. The biological processes shown in this paper are exhaustive, that is, we tried to avoid nested processes within other more general.

### Western blot analysis

The results validation was done through OXPHOS proteins immunodetection (Wb) of OXPHOS proteins and bioenergetic signature. Mitochondrial extracts were obtained from 6 million T98G and U87MG cells. These were subjected to SDS-PAGE 12% system described in Laemelli (1970) [31]. Gels ran for 2h at 100V. Proteins separated by SDS-PAGE were transferred to PVDF membrane, as described in Towbin (1979) [32], at 100V for 1h; an antibody against a subunit of each OXPHOS complex: NDUFA10, CI (1:2000); subunit 70 kDa, CII (1: 10000); core 2, CIII (1:4000); subunit IV, CIV (1:1000) and beta subunit ATP synthase or CV(1:1000) was tested. The reaction bands were detected by chemiluminescent (Millipore, WBKLS0500) and read on to C-Digit Blot Scanner (LI-COR).

## Results

### Spot selection by Random Sampling and Principal Component Analysis

Three of the 400 spots selected across all gels surface, regardless size, intensity or difference in abundance between cell lines, did not pass the quality control; 161 spots were specific for T98G or U87MG. Therefore, the PCA was applied to 236 spots shared by both lines (Supplementary Table 1). According to the PCA, the total variation in the spots abundance in all gels can be explained by five principal components (Fig. 1A). The first component (PC1) holds 63% of whole explained variance (Fig 1B) whiles the others four components together explain only the 37% of the variance remaining. PC1 also distinguish between T98G and U87MG gels (Fig. 1C). 165 spots from CP1 were selected according to its relative importance between components (square cosine of the correlation matrix between the components and the spots (Supplementary Table 2)) to MALDI-TOF identification, 114 of them show a positive correlation and 51 a negative one (Supplementary Table 3). The first set of spots showed more mean abundance in T98G and smaller in U87MG, different than the second set, which is more abundant in U87MG (Fig. 1D). 20 specific spots in T98G and 20 in U87MG (randomly selected too) were added.

**Fig. 1.**
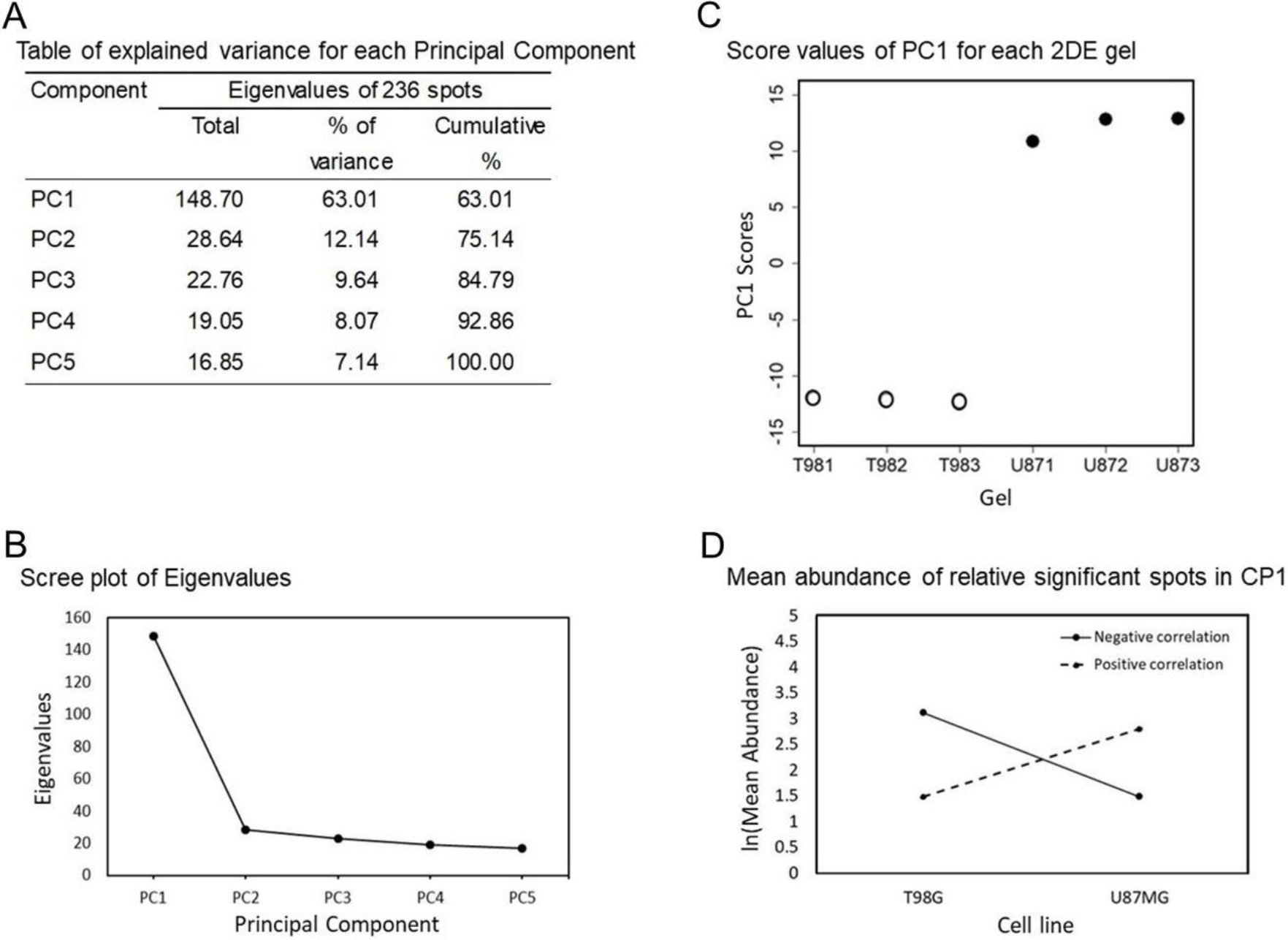
PCA analysis was done on spot abundance to get 5 PCs explaining the total variation in the data (A). As PC1 explains 63% of whole variance and is located to the left of inflection point in the Scree plot line (B). Thus, PC1 scores for al gels (C) distinguishes between experimental groups. Finally, average abundance profile plot (D) shows the behavior of significant spots in PC1, with positive correlation between spots and PC1 (dot line), others with negative one (solid line); suggesting a molecular transition.

### T98G and U87MG landscapes

As a result of Random Sampling and the PCA 89 identified spots (Supplementary Table 4) had a homogeneous distribution in T98G and U87MG gels (Figs. 2A and 2B), unrelated to size, intensity or difference in abundance between cell lines, assuring the representativeness of whole mitochondrial proteome in these lines.

**Fig. 2.**
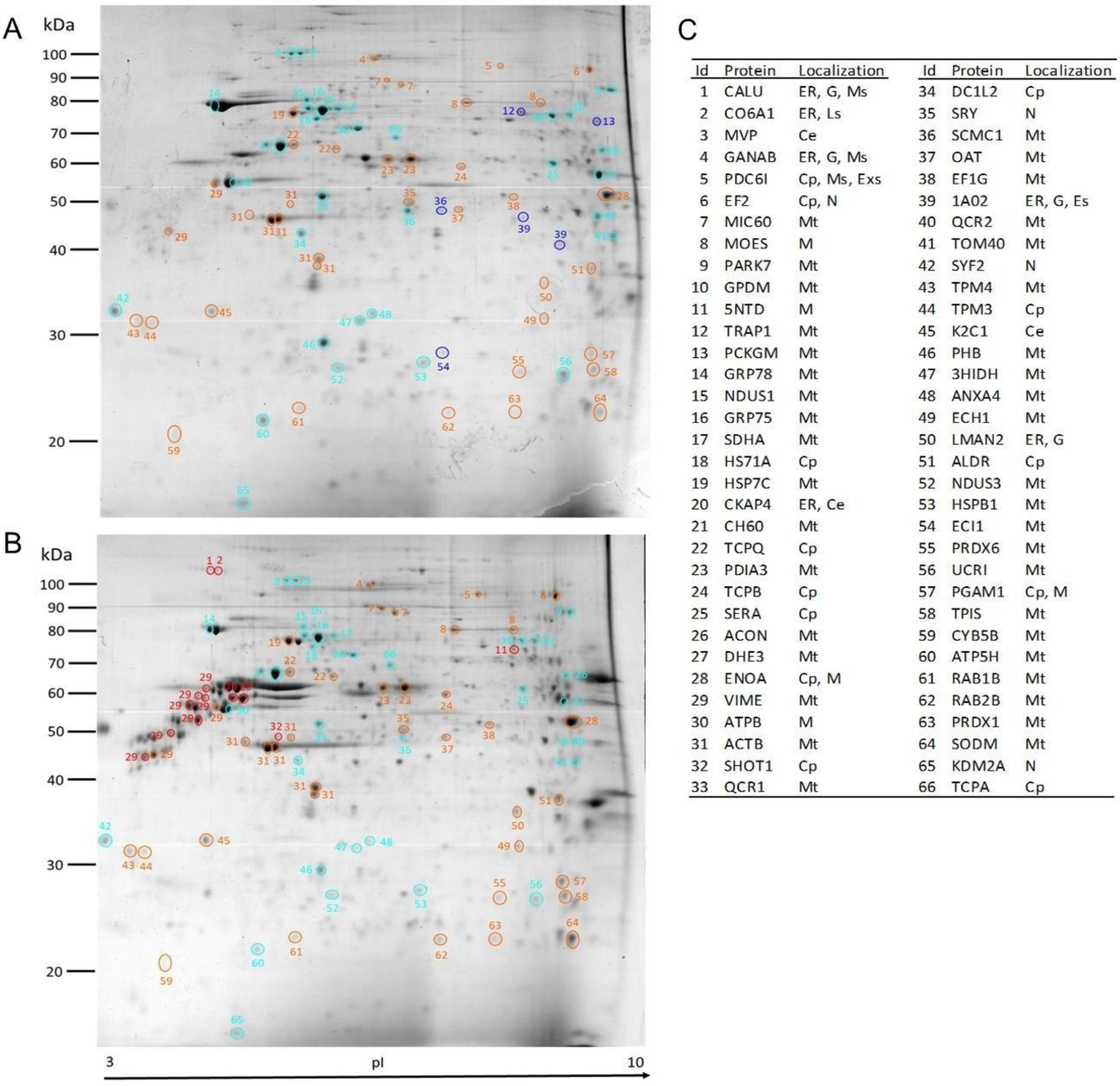
Mitochondrial proteome distribution in T98G (A) and U87MG (B) gels of identified proteins, and their cell localization (C) as result of random sampling and PCA. Circles in blue, blue light, red and orange are specific and overexpressed proteins in T98G and specific and overexpressed proteins in U87MG respectively. Cell localization was obtained according MitoMiner database and GeneCard Suite. Cytoskeleton (Ce), Cytoplasm (Cp), Endoplasmic Reticulum (ER), Endosome (Es), Exosome(Exs), Golgi Apparatus (G), Lysosome (Ls), Melanosome (Ms), Membrane (M), Mitochondria (Mt) and Nucleus (N).

Since mitochondria are multifunctional organelles, proteins with different origin can colocalize in them. The identified protein dataset was compared against MitoMiner database to recognize mitochondrial proteins. We found that T98G show more mitochondrial proteins (72%, 24 proteins) than U87MG (44%,15 proteins). “Foreign” proteins were located in Cytoplasm, Plasmatic Membrane, Endoplasmic Reticulum, Golgi, and Nucleus according to GeneCards Suite [33] (Fig. 2C).

The initial mitochondrial proteome PPI networks were built with 66 proteins represented by the 89 identified entities. In T98G, 33 proteins (29 proteins more overexpressed and 4 specifics). U87MG initial PPI network groups 33 proteins (28 more abundant and 5 own) (circles in Figs. 3A and 3B).

**Fig. 3.**
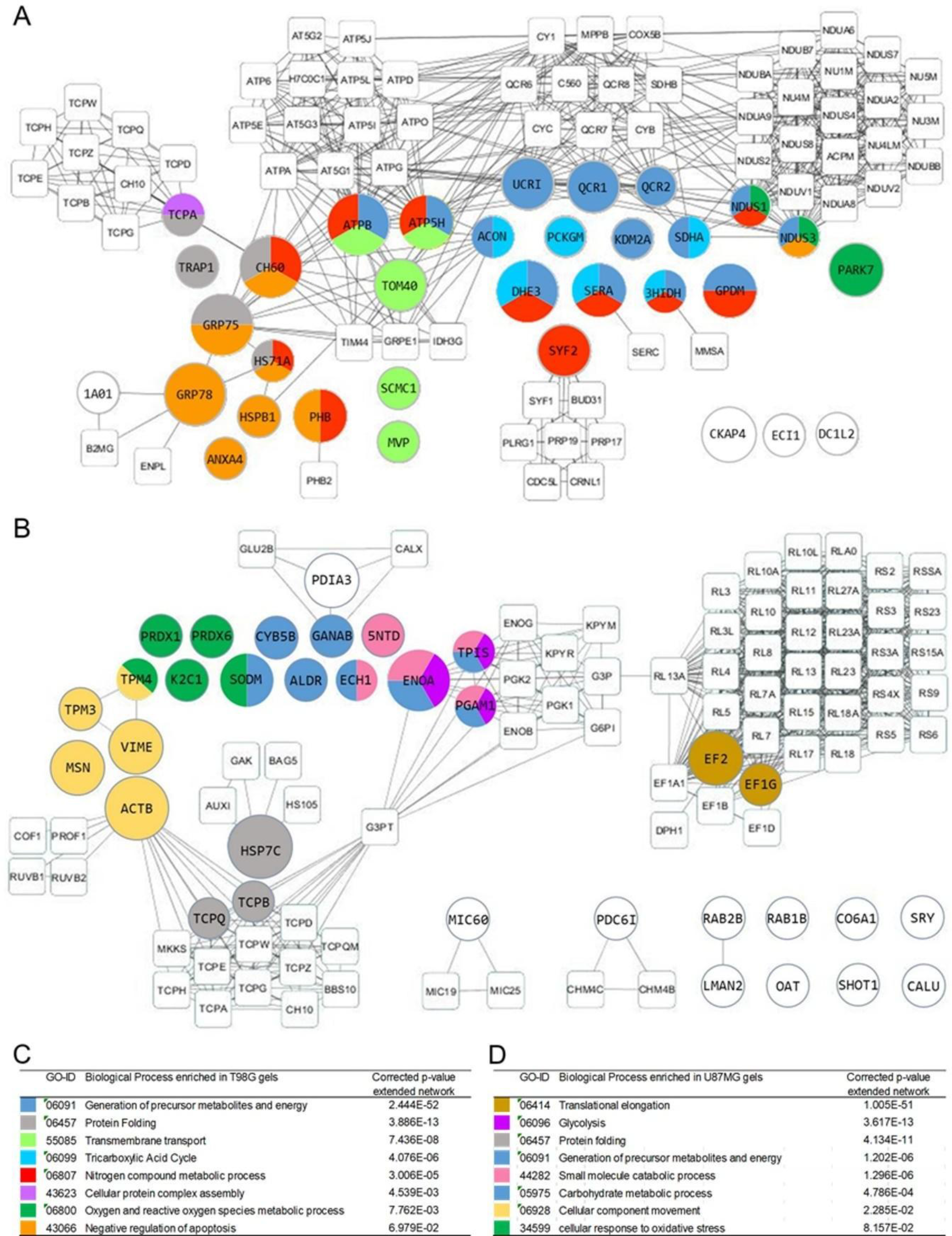
The biological processes of each glioblastoma cell line is shown by colors. The processes were obtained by protein aggregation (squares) from T98G (A) and U87MG (B) initial PPI networks (circles). The size of the circles (small, medium and large) represents the relative abundance (low, regular and high, respectively) of proteins.

To get a better landscape of mitochondrial function in each cell line, initial PPI networks were extended (Squares in Figs. 3A and 3B) resulting in a T98G extended PPI network (Fig. 3A) where mitochondrial processes dominant showing functional mitochondria. One of the best-represented processes here is the “Generation of precursor metabolites and energy” process (p-value 2.4E-52, Fig. 3C) with OXPHOS (UCRI, QCR1, QCR2, NUDS1 and NUDS3) and ATPB y ATP5H proteins included, coupled with the “Oxygen and reactive oxygen species metabolic process” (p-value 7.6E-03, Fig. 3C) and “Tricarboxylic Acid Cycle” (p-value 4.1E-06, Fig. 3C) represented by ACON, SDHA, DHE3, SERA and 3HIDH. Another of the process present in the T98G network is the “Nitrogen compound metabolic process” (p-value 3.0E-05, Fig. 3C), which connects with OXPHOS trough “Transmembrane transport” process (p-value 7.4E-08, Fig. 3C) and CH60 and HS71A proteins. This process is also chained with “Negative regulation of apoptosis” (p-value 6.9E-02, Fig. 3C) and “Protein folding” (p-value 3.9E-13, Fig. 3C) proteins. It is interesting that other cellular process without enough statistical representation let see different typical mitochondrial pathways like ◻-oxidation, represented by ECI1 (Fig. 3C). U87MG extended PPI network (Fig. 3B) shows many differences with that of T98G. Although this net is more fractionated, it is possible to recognize some enriched biological process. One of the more remarkable, found in carcinogenesis, is the change on energetic metabolism, represented by “Glycolysis” (p-value 3.6E-13, Fig. 3D) proteins ENOA, PGAM1 and TPIS, “Generation of precursor metabolites and energy” (p-value 1.2E-06, Fig. 3D)” and “Small molecule catabolic process” (p-value 1.3E-06, Fig. 3D).

“Cellular component movement” (p-value 2.3E-02, Fig. 3D) and Cellular response to oxidative stress” (p-value 8.1E-02, Fig. 3D) processes, related to cell proliferation and invasive cell capacity, were found in U87MG. Also, it were found “Translational elongation” process (p-value 1.0E-51, Fig. 3D), that groups EF2 y EF1G proteins and “Protein folding”, (p-value 4.1E-11, Fig. 3D), which are related to an increased protein translation for augment biomass since many of them are molecular chaperones (HSP7C, TCPB, TPCQ). This U87MG landscape shows mitochondria with modified cellular and metabolic functions and many interactions with ER and Golgi body (Figs. 2C and 3B), suggesting that mitochondria readjust its cellular process according to carcinogenesis needs.

To know if any biological process is grouping the most abundant proteins, a k-means analysis was performed. The proteins were classified, in function of its relative abundance, as little, regular or very abundant (small, medium and large circles respectively in Figs. 3A and 3B). We found low, regular and very abundant proteins in all biological processes in both cell lines; showing no correlation between the overrepresentation of some biological process and the abundance of the proteins that represent it.

### Protein and biological process validation for mitochondrial proteomic signature

Due to “Generation of precursor metabolites and energy process” was one of the best-represented processes in T98G (p-value 2.4E-52) and U87MG (p-value 1.2E-06), the OXPHOS protein expression (I-IV complexes plus ATP synthase) was verified on both cell lines by WB assays. A clear diminished OXPHOS system expression in U87MG cells was found (Fig. 4A). Also, as “Glycolysis” was the most enhanced process in U87MG (3.6E-13), the bioenergetic signature was assayed too (Fig. 4B). The results support OXPHOS expression finding since U87MG cells expressed more glycolytic proteins comparing to T98G, where OXPHOS system is dominating. This result confirmed PPI networks built on the basis to random sampling and PCA analysis.

**Fig. 4.**
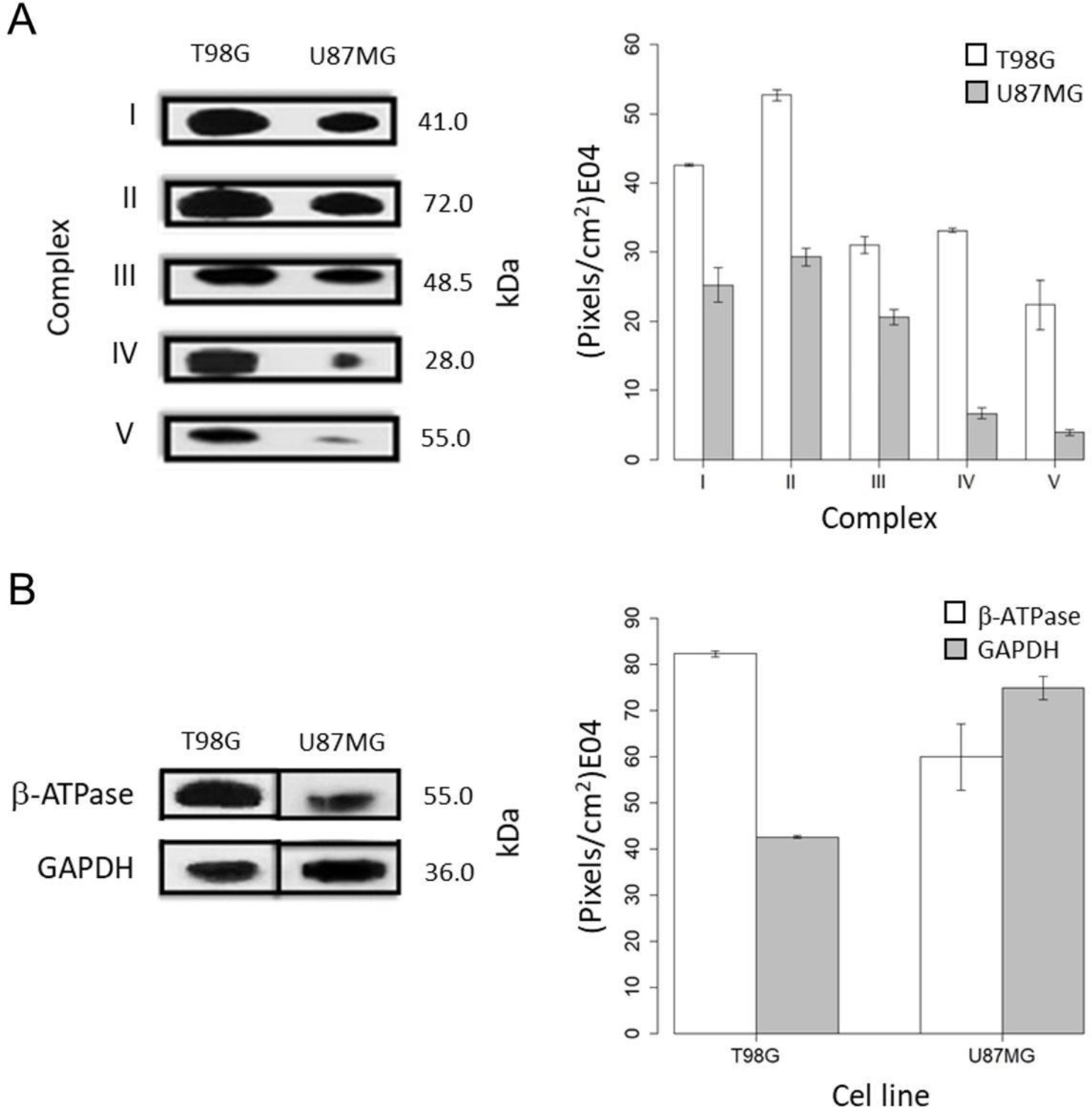
OXPHOS validation (A) and bioenergetic signature (B). To the left in figure the expression bands obtained by western blot are observed. To the right is plotted the average and 95% average confidence interval of density bands, calculated by triplicate for each complex, β-ATPase and GAPDH.

**Fig. 5.**
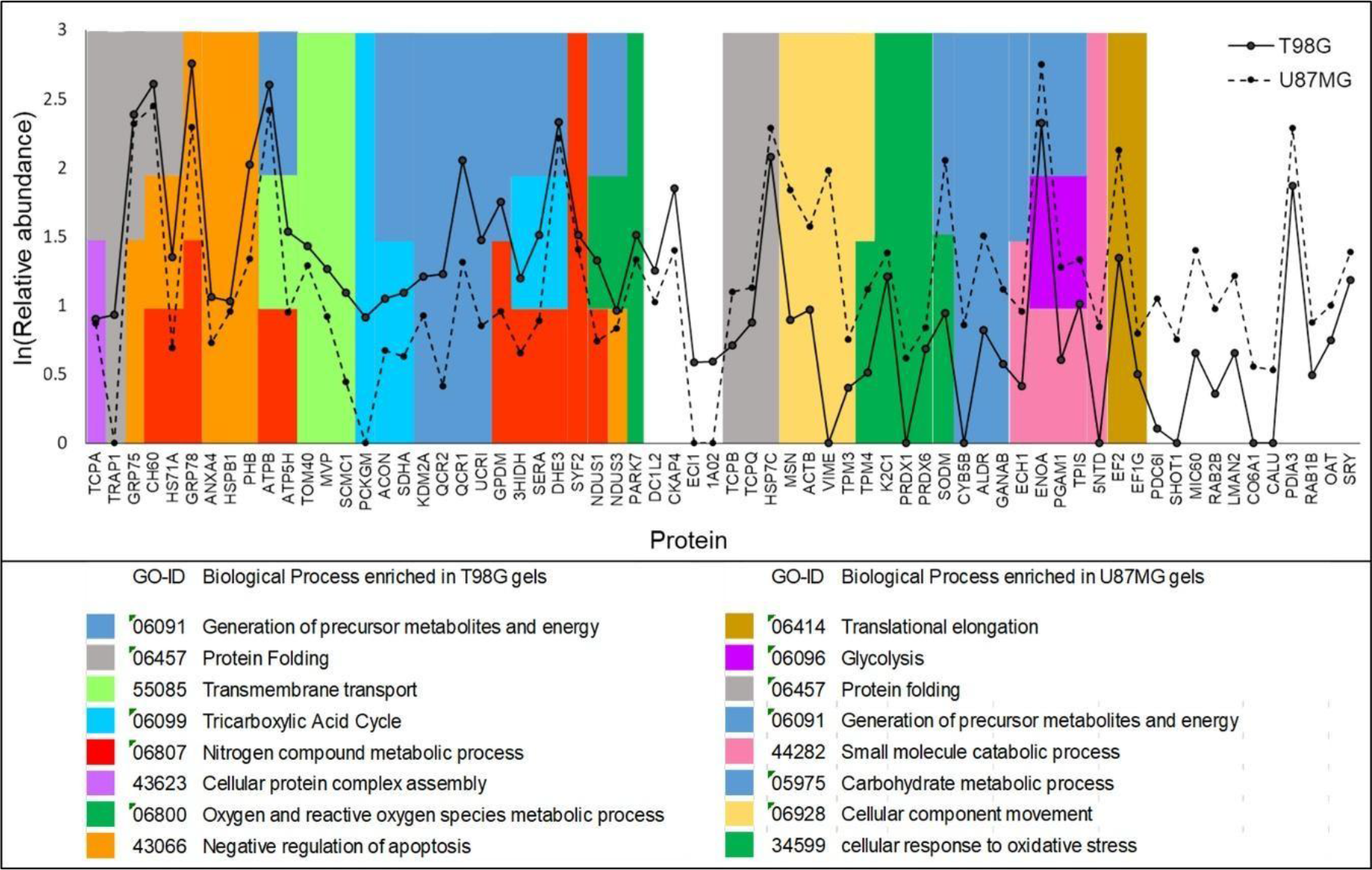
Proteomic signature generated with the mean abundance profile of the three replicates of T98G (solid line) and U87MG (dot line). Colors locate the proteins in the main biological processes identified for each cell line.

## Discussion

Besides ATP synthesis mitochondrion is a multifunction organelle, which is involved in many cellular processes. Mitochondria proteome is versatile and reacts to different cellular conditions; many complex diseases including cancer show a mitochondrial roll. The best-known mitochondrial change in cancer is Warburg effect: an energetic metabolism shift to glycolysis, as a mean energy source, instead of mitochondrial OXPHOS. It used to be believed that Cancer cells were related to mitochondrial dysfunction. However, in some cancer types, exist enough evidence showing complete functional mitochondria, able to follow cellular transformation[15]. Nowadays, it has believed that mitochondria follow cancer development sensing and regulating different molecular signals [34–36]. Thus, there are many mitochondrial proteins or mitochondrial processes that could be clinical targets or biomarkers.

Our results describe two well-differentiated states from mitochondrial proteome data. Our analysis by RS on 2D SDS PAGE and spot abundance by PCA, allow the detection of mitochondrial and cellular pathways distinguished. The data are in resonance with biochemical and proteomic evidence [12,13]. Our approach renders a landscape close to molecular cancer dynamics according to published evidence on glioblastoma biology and systematics [15,37–39], enabling to raise a proteomic signature for T98G and U87MG glioblastoma cells with the best representative biological process according to each mitochondrial proteome. PCA raise five components, the first component explains 63 % of total variation in spot abundance data and have proteins that distinguish between T98G and U87MG cells. PC1 could be renamed as “Energy metabolism shift” since it represents many of the processes involved in the Warburg effect.

Random spot selection and PCA from direct experimental data before identification point out a specific PPI network for T98G and U87MG cells, where the energetic metabolic shift to glycolysis as the mean ATP source occurs. U87MG cells represent an advanced, invasive and malignant cancer state vs T98G cells, which represent an earlier state with OXPHOS metabolism. According to our PPIs, T98G cells show typical mitochondrial functions (OXPHOS, TCA, lipid metabolism, etc.) but also another more cancer-related process (Apoptosis evasion, proliferation with SYF2, HSPA8, amino acids metabolism with (GLUD1, 3HIDH); or chaperon response. On the other hand, U87MG cells show promiscuous interactions with ER and Nucleus, a maintained chaperone response and DNA translation to proteins, a very advanced state with invasion-related proteins.

This characterization could define different cancer state or intervals and works for other cancer types too. On this way, T98G cells could represent an earlier cancer state with a molecular landscape similar to “oxidative tumors”, where ATP comes from OXPHOS system fueled by lipid (ECl1, GPD2) and amino acids metabolisms, like glutamine (GLUD1), as it has been observed in some glioblastoma cases [15,40]. U87MG shows a very different state, in which glycolysis is well represented (ENOA, TPIS, PGAM1), supposing an enhanced Warburg effect. Also, oxidative stress response as is has been reported [13,41] and many non-mitochondrial but close cancer-related proteins reported in advanced tumors [42]. U87MG mitochondria show mobility or migration proteins related to the cytoskeleton (VIME, ACTB, MSN, TPM3), and vesicle formation (RAB1B, RAB2B, LMAN2). Another less frequent processes were well represented such as DNA translation (EF2, EF1G) into proteins; this increase could be associated to biomass increase or metabolic energy source, since many proteins folding chaperones (TCPQ, TCPB) were observed.

Our procedure for protein analysis enables us to determine various simultaneous cell processes besides metabolic shifting. A remarkable glioblastoma molecular feature is the chaperones response, where some biomarkers [43] or therapy targets [44] could be found. Here are presented TRAP1 (HSP90 homologous), GRP78, GRP75 and HSPB1 proteins able to regulate some mitochondrial metabolic pathways and stabilize cancer cells through apoptosis evasion [45]; or could be involved in drug surveillance [38].

The finding of non-mitochondrial proteins in our study is not a surprise. Basal mitochondria function includes the interaction with other cell organelles, mainly ER and the Nucleus. Our data show some nuclear (SRY) or ER (CALU, CO6A1) proteins. In U87MG cells there are more interactions between these proteins, suggesting a specificity of these interactions on advanced cancer. This landscape resembles autophagy, a central process in advanced states of cancer, which enable cancer cell surveillance because of the recycling of metabolites and nutrients [46,47]. In addition, there are proteins for amino-acids and purines metabolism making possible the phagosomes formation [48]. Autophagia renders biomass bricks or stress response molecules synthesis (i.e., amino acids generation by proteolysis, recycling and protein synthesis for fueling other pathways (TCA)), when basal or other metabolites are not available (Formation of metabolic precursors, RAB).

With this information, a proteomic signature for T98G and another for U87MG was proposed, defining concrete cell processes and temporality. Unlike other protein signatures which look for more straight aims, like biomarkers search using other biological models (plasma or cerebrospinal liquid proteins) where proteins surpassing significant abundance changes and overseeing some cellular process, resulting in inadequate descriptions [49,50].

## Conclusions

The random sampling of spots and their abundance PCA before protein identification are tools that allow us to see a fine landscape of the most relevant biological process or functions in each cell type or glioma carcinogenesis state; with this information we are able to build a representative mitochondrial proteomic signature specific for T98G and U87MG glioblastoma cell lines, where overrepresented biological processes are highlighted with whole mitochondrial proteins identified. This signature shows a clear difference between two glioblastoma stages, one with mitochondrial type (OXPHOS) metabolism and, the other, a glycolytic, more aggressive, invasive and metastatic cancer type.

Our data match with the notion of mitochondria as a dynamic organelle following and sensing the molecular events taking place during carcinogenesis. Through this close relationship is possible to take a temporal picture of cancer stages or types. It also shows that a well-selected spot sample and a correct data analysis of mitochondrial proteome can define the biological events succeeding in cellular transformation. Thus, the notion that T98G could represent an earlier glioblastoma state bring the opportunity to focus in an earlier cancer-related events, such as apoptosis evasion, and target the chaperone system as a therapeutic diana to avoid cancer development.

## Acknowledgments

We are grateful to Dr. Hector Vargas for insightful discussion. This paper constitutes a partial fulfillment of the Post-Graduate Program in Biological Sciences of the Universidad Nacional Autónoma de México (UNAM). Leopoldo Gómez-Caudillo acknowledges to Consejo Nacional de Ciencia y Tecnología (CONACyT) for the scholarship No. 375815, and to Instituto Mexicano del Seguro Social (IMSS) (FIS/IMSS/PROT/G13/1206). This work was supported in part by Consejo Nacional de Ciencia y Tecnologíia (CONACyT) Grant 220790 and Directión General de Asuntos del Personal Académico-Programa de Apoyo a Proyectos de Investigatión e Inovación Tecnológica, Grant IN213216.

